# CMTr cap-adjacent 2’-*O*-ribose mRNA methyltransferases are required for reward learning and mRNA localization to synapses

**DOI:** 10.1101/2021.06.24.449724

**Authors:** Irmgard U. Haussmann, Yanying Wu, Mohanakarthik P. Nallasivan, Nathan Archer, Zsuzsanna Bodi, Daniel Hebenstreit, Scott Waddell, Rupert Fray, Matthias Soller

## Abstract

Cap-adjacent nucleotides of animal, protist and viral mRNAs can be dynamically *O*-methylated at the 2’ position of the ribose (cOMe). The functions of cOMe in animals, however, remain unknown. Here we show that the two cap methyltransferases (CMTr1 and CMTr2) of *Drosophila* can methylate the ribose of the first nucleotide in mRNA. Double-mutant flies lack cOMe but are viable. Consistent with prominent neuronal expression, they have a reward learning defect that can be rescued by conditional expression in mushroom body neurons before training. Among CMTr targets are cell adhesion and signaling molecules relevant for learning and cOMe is required for local translation of mRNAs at synapses. Hence, our study reveals a mechanism to co-transcriptionally prime mRNAs by cOMe for localized protein synthesis at synapses.

## Introduction

Methylation of cap-adjacent or internal nucleotides in messenger RNA (mRNA) is a major post-transcriptional mechanism to regulate gene expression. Methylation of mRNA is particular prominent in the brain, but the molecular function of methylated nucleotides and their biological roles are poorly understood ^1-5^.

Methylation of cap-adjacent nucleotides is an abundant modification of animal, protist and viral mRNAs, that varies in different tissues and transcripts ^6-17^. Dynamic *O*-methylation at the 2’ position of the ribose (cOMe) of cap-adjacent nucleotides is introduced co-transcriptionally by two dedicated cap methyltransferases (CMTr1 and CMTr2) after capping at the beginning of an mRNA to a characteristic 5’-5’ linked *N7*-methylated guanosine ^18-20^.

The main function of the cap is to protect mRNAs from degradation and to recruit translation initiation factors, but also to promote splicing and 3’ end processing ^21^. The cap is initially bound in the nucleus by the cap binding complex (CBC), consisting of CBP20 and CBP80. Upon export from the nucleus, CBC is replaced by eIF4E, which is predominantly cytoplasmic and rate-limiting for translation initiation ^22,23^. *N7*-methylation of the cap guanosine is critical for both CBC and eIF4E binding. The importance of cap-adjacent nucleotide methylation in animal gene expression, however, remains elusive, but is known to be essential in trypanosomes and viruses including SARS-CoV-2 for propagation ^15^.

## Results

### *CMTrs* act redundantly

To elucidate the biological function of cap-adjacent 2’-*O*-ribose methylation (cOMe) in animals we made null mutants of the *CMTr1* (*CG6379*) and *CMTr2* (*adrift*) genes in *Drosophila*. We generated small intragenic deletions in each gene by imprecise excision of a *P*-element transposon to make *CMTr1*^*13A*^ and *CMTr2*^*M32*^ mutant flies (**Fig. 1a-c**). Both of these genetic lesions remove the catalytic methyltransferase domain from the encoded CMTr1 and CMTr2 protein. Perhaps surprising, these mutant flies are viable and fertile as single and double mutants, exhibiting a slightly reduced survival to adulthood after hatching from the egg (**Fig. 1d**) and climbing activity in negative geotaxis assays (**Fig. 1e**).

**Figure 1.**
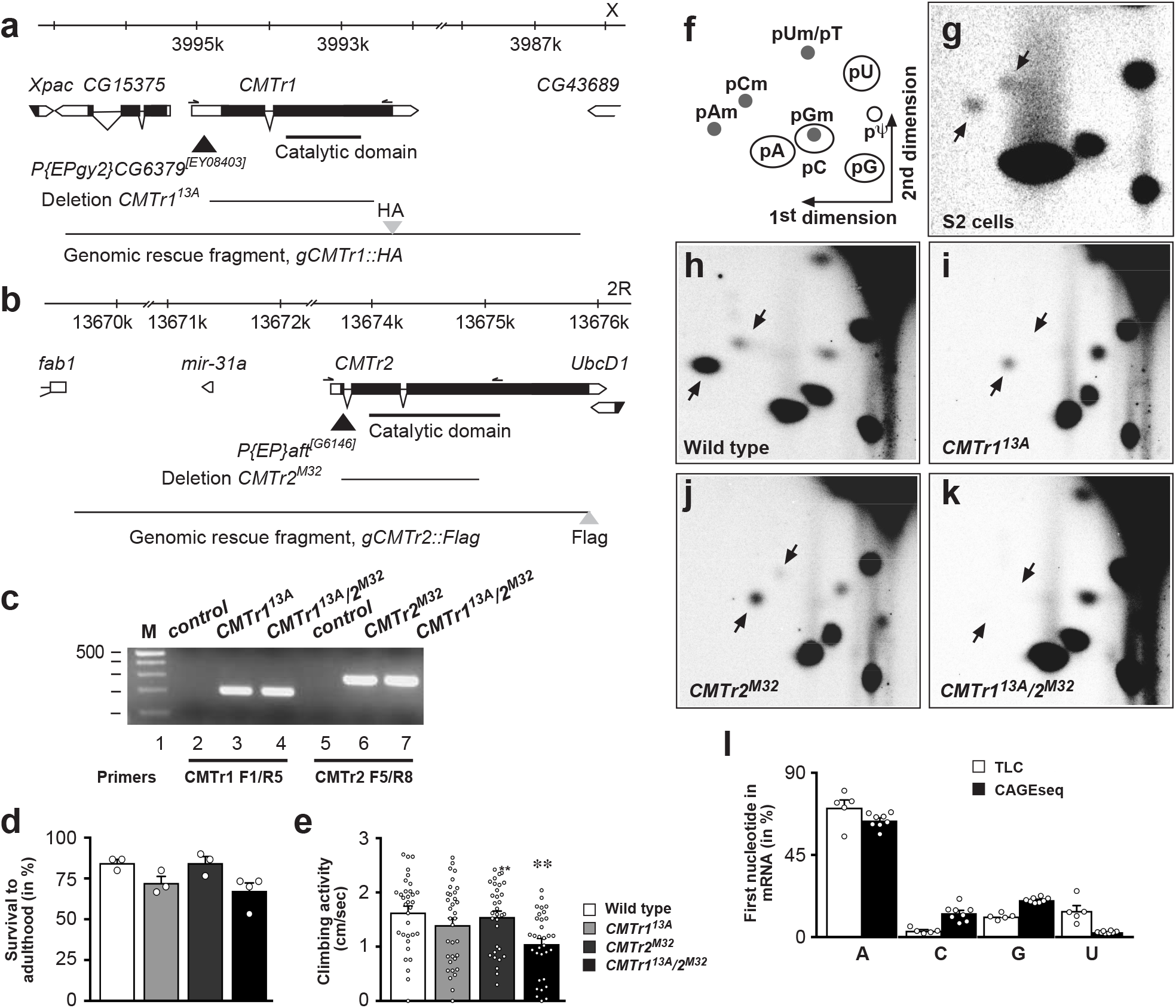
Analysis of *CMTr1* and *CMTr2* null mutants and mRNA cap 2’-*O*-ribose methylation in *Drosophila*. **(a and b)** Genomic organization of the *CMTr1* and *CMTr2* loci depicting the transposons (black triangle) used to generate the deletions *13A* and *M32*, which are null alleles. Genomic rescue fragments tagged either with hemaglutinin (HA, a) or FLAG (b) epitopes are indicated at the bottom. **(c)** Validation of *CMTr1*^*13A*^ and *CMTr*^*M32*^ single and double mutants by genomic PCR. **(d)** Survival of flies to adulthood after hatching from the eggshell (n=3-4). **(e)** Climbing activity assessed by negative geotaxis assays, n=40, p≤0.001. **(f)** Schematic diagram of a 2D thin layer chromatography (TLC) depicting standard and 2’-*O*-ribose methylated nucleotides. **(g-k)** TLCs showing modifications of the first cap-adjacent nucleotides of S2 cells (e), adult control (f) and *CMTr1*^*13A*^ and *CMTr2*^*M32*^ single (g, h) and double (i) mutant females. **(l)** Quantification of the mRNA first nucleotide from TLC (n=5) and CAGEseq data (n=8) from adult *Drosophila* and S2 cells, respectively.

We next detected cOMe in purified mRNAs from S2 cells and adult female flies by thin layer chromatography (TLC) (**Fig. 1f**). In S2 cells, we detected cOMe on adenosine (pAm) and cytosine (pCm, **Fig. 1f and g**), but in female flies predominantly pAm was present (**Fig. 1h**). Although, single mutants in CMTr1 or CMTr2 still had cOMe, the double mutants were devoid of cOMe suggesting that these two enzymes have overlapping function and are both able to methylate the 2’-*O*-ribose of the first transcribed nucleotide (**Fig. 1i-k**).

Moreover, our TLC analysis of the first nucleotide in *Drosophila* mRNAs shows a strong preference for A (**Fig. 1l**), which is consistent with the transcription initiator motif (Inr) sequence YYANWYY (Y: pyrimidine, N: any nucleotide and W: A or T) obtained from *Drosophila* by CAGEseq (**Fig. 1l**) ^24^.

### *CMTrs* are broadly expressed

Global expression studies of *CMTr1* and *CMTr2* showed that both are expressed throughout development in a broad range of tissues with elevated *CMTr1* levels during early embryogenesis and a peak of both in prepupae (**Supplementary Fig. 1a and b**)^25,26^. Both *CMTr1* and *CMTr2* show higher expression in larval brains and to some extent in the adult nervous system and in ovaries (**Supplementary Fig. 1b**). *CMTr2* is also highly expressed in testis and trachea, which is consistent with a previously described transient role in tracheal development ^27^.

Analysis of expression from epitope-tagged genomic rescue constructs in the larval ventral nerve cord and adult brains revealed expression of both CMTr1 and 2 primarily in a pan-neural pattern with a predominantly nuclear localization of both as compared to the nuclear neuronal marker ELAV (**Supplementary Fig. 1c-n**). To obtain a clearer view of the intracellular localization we stained epitope tagged CMTr1 and CMTr2 in third instar salivary glands (**Supplementary Fig. 1o-w**). CMTr1, and to lesser extent CMTr2, were both enriched in the nucleus, but excluded from the nucleolus. There was also prominent localization of CMTr2 to the cytoplasm and the cell membrane. In addition to cytoplasmic staining, CMTr2 also prominently localizes to the cell membrane, and this is also somewhat evident for CMTr1.

### Reward learning requires cap methylation

mRNA modifications have been associated with neurological disorders and intellectual disabilities in humans ^4,28^. Given the increased expression of CMTrs in the brain, we tested CMTr mutant flies for learning and memory using appetitive conditioning that rapidly forms protein-synthesis dependent memory ^29^.

Immediate (3 min) and 24 h memory of single *CMTr1*^*13A*^ and *CMTr2*^*M32*^ mutant flies was indistinguishable from that of wild-type controls. However, both immediate and 24 h memory were significantly impaired in *CMTr1*^*13A*^; *CMTr2*^*M32*^ double mutant flies (**Fig. 2a and b**). These memory performance deficits were restored by introducing transgenes encoding genomic fragments for both *CMTr1* and *CMTr2* (**Fig. 2c**), indicating that the learning deficits arise from the absence of cOMe function.

**Figure 2.**
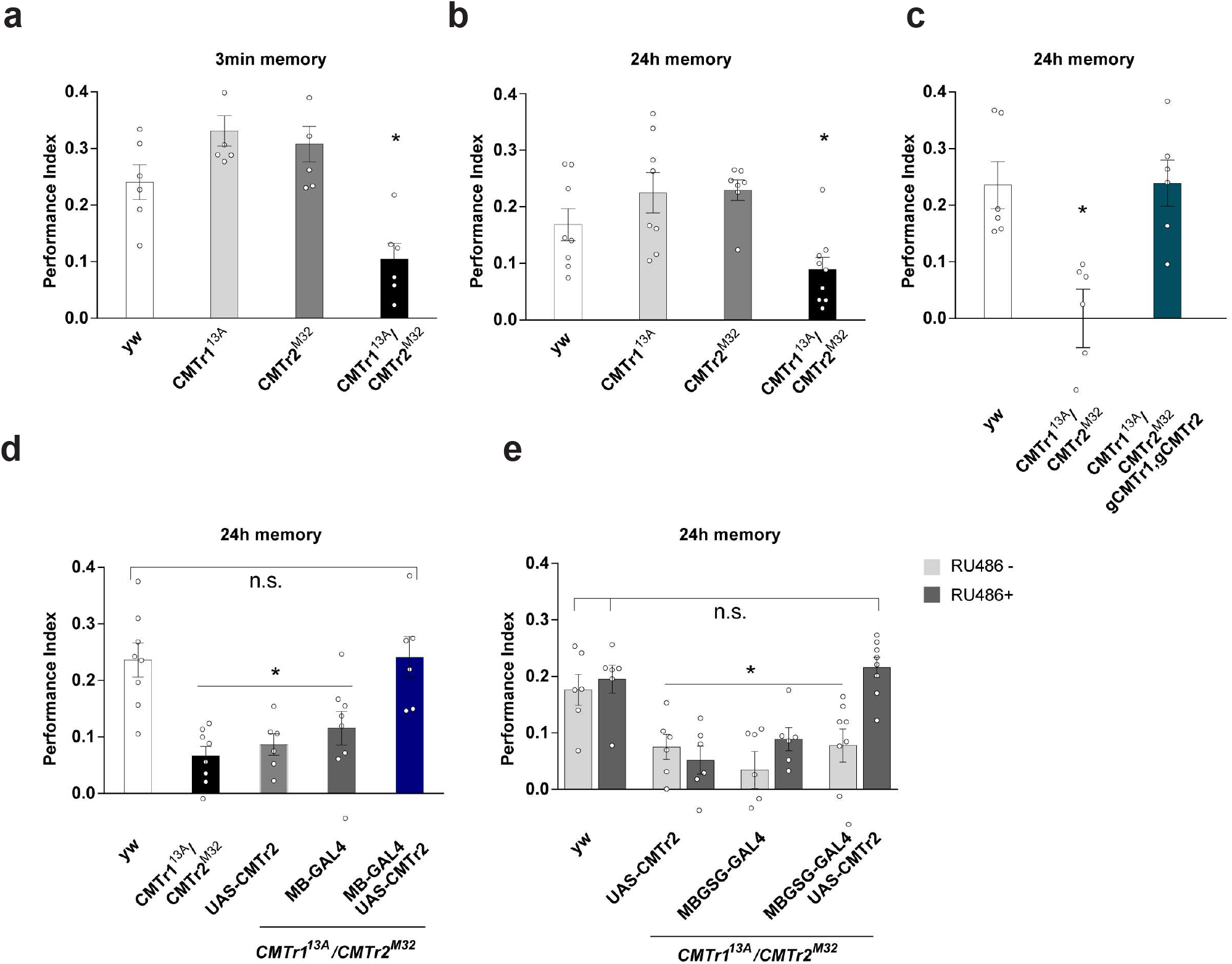
mRNA cap 2’-*O*-ribose methylation is required for reward learning in *Drosophila*. **(a and b)** Appetitive memory immediately (a) and 24 hour (b) after training of control and *CMTr1*^*13A*^ and *CMTr*^*M32*^ single and double mutant flies shown as mean±SE. n=8 for A and n=6 for B, p≤0.006. **(c)** Rescue of the learning defect in *CMTr1*^*13A*^; *CMTr2*^*M32*^ double mutant flies by genomic fragments shown as mean±SE. n=6, p=0.002. **(d and e)** Rescue of the learning defect in *CMTr1*^*13A*^; *CMTr2*^*M32*^ double mutant flies by constitutive (d) or conditional (e) expression of CMTr2 in mushroom bodies from *UAS* shown as mean±SE. n=6, p≤0.0001.

We also tested performance of *CMTr1*^*13A*^; *CMTr2*^*M32*^ mutant flies using aversive olfactory conditioning which pairs one of two odors with an electric shock. Surprisingly, aversive learning of *CMTr1*^*13A*^; *CMTr2*^*M32*^ double mutant flies was indistinguishable from that of control flies, which suggests specificity for the reward learning defect (**Supplementary Fig. 2a**). Moreover, mutant flies behave normally when exposed to the repellent odors and they can detect sugar (**Supplementary Fig. 2b and c**). These sensory controls and the wild-type aversive learning performance of *CMTr1*^*13A*^; *CMTr2*^*M32*^ also suggest that cOMe deficiency somehow specifically impairs reward learning.

Olfactory learning and memory in *Drosophila* is coded within the neuronal network of the mushroom bodies (MBs)^30^. Valence learning can be coded as changes in the efficacy of synaptic junctions between odor-activated Kenyon Cells (KCs, the intrinsic cells of the MB) and specific mushroom body output neurons. We therefore tested whether the reward learning defect of *CMTr1*^*13A*^; *CMTr2*^*M32*^ mutant flies could be rescued by restoring cOMe expression to KCs. Expressing a *UAS-CMTr2* transgene in the KCs using *MB247-GAL4* rescued the learning deficits of *CMTr*^*13A*^; *CMTr*^*M32*^ double mutant flies (**Fig. 2d**).

Next, we investigated whether the reward learning phenotype of *CMTr*^*13A*^; *CMTr*^*M32*^ double mutant flies arose from a developmental origin, or from loss of an acute function in the adult stage. The gross morphology of the adult MBs appears to be normal in *CMTr1*^*13A*^; *CMTr2*^*M32*^ mutants as judged from expressing a *UAS-EGFP* transgene with *MB247-GAL4*, or with the KC-subtype restricted drivers *NP7175-GAL4* (αβ core KCs), *0770-GAL4* (αβ surface KCs) or *1471-GAL4* (γ KCs, **Supplementary Fig. 3a**). Interestingly, restoration of *CMTr2* expression to these more restricted KC subsets did not rescue the learning defect of *CMTr*^*13A*^; *CMTr*^*M32*^ double mutant flies (**Supplementary Fig. 3b**).

We next tested whether the reward learning defect of *CMTr*^*13A*^; *CMTr*^*M32*^ double mutant flies could be rescued by inducing CMTr2 expression just before training in adult flies. Since *MB247-GAL4* was able to restore learning, we employed a *MB247*-driven Gene-Switch (GS) to conditionally induce CMTr2 expression by feeding flies with RU486. Only *CMTr*^*13A*^; *CMTr*^*M32*^ flies that also harboured the *MB247-GS* and *UAS-CMTr2* transgenes exhibited restoration of memory performance when fed with RU486 (**Fig. 2e**). Together these experiments suggest that cOMe in the MB KCs plays a key role in olfactory reward learning.

### *CMTr* loss increases transcript abundance

To investigate the impact of cOME on gene expression, we performed RNA sequencing on cOMe deficient and control flies. Differential gene expression analysis revealed 197 and 701 genes that were significantly down- and up-regulated in *CMTr*^*13A*^; *CMTr*^*M32*^ double mutant flies as compared to wild type controls (adjusted p-value<0.05, at least twofold change, **Fig. 3a, Data S1**). GO term analysis revealed significant up-regulation of genes involved in metabolism, receptor signalling and cell adhesion (**Data S2**). To obtain a high confidence list of significantly differentially regulated genes, we took genes threefold differentially regulated (80 and 244 genes down- and up-regulated in double mutant flies compared to controls) and analysed them according to gene function by annotated protein domains. This analysis confirms prominent effects on gene networks involved in metabolism, cellular signaling and structural cell components, including a number of cell adhesion molecules and is qualitatively different from loss of m^6^A or by regulation of synapse numbers by the transcription factor *erect wing* (**Fig. 3b, Data S2**)^31-33^.

**Figure 3.**
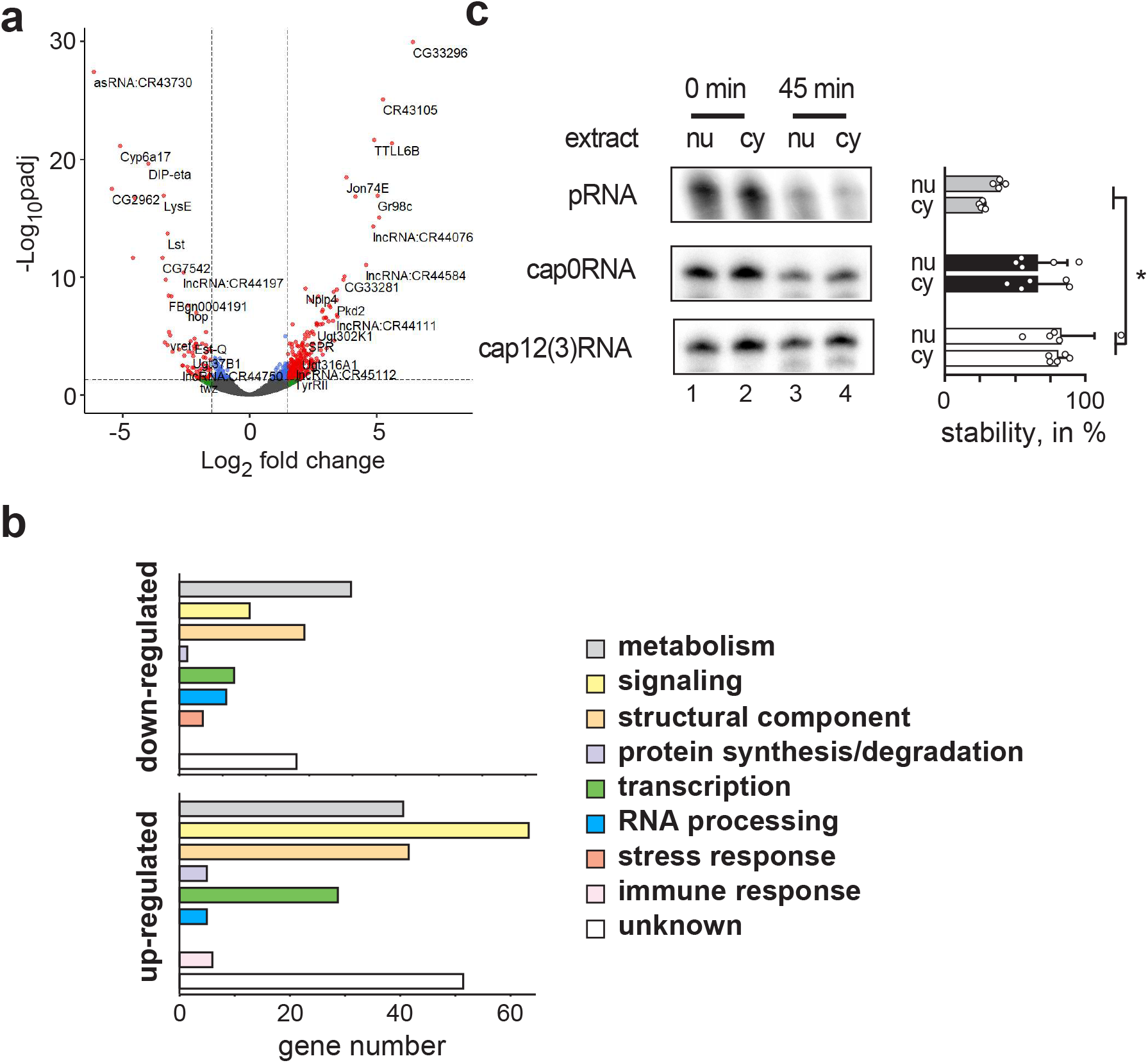
Impact of mRNA cap-adjacent 2’-*O*-ribose methylation on gene expression and RNA stability. **(a)** Volcano plot depicting differentially expressed genes in *CMTr1*^*13A*^; *CMTr2*^*M32*^ double mutant flies compared to control flies. **(b)** Functional classification of up-(bottom) and down-regulated (top) genes in *CMTr1*^*13A*^; *CMTr2*^*M32*^ double mutant flies compared to control flies. **(c)** Incubation of monophosphorylated RNA (pRNA) and capped RNA with or without 2’-*O*-ribose methylation of cap-adjacent nucleotides in nuclear (nu) and cytoplasmic (cy) extracts from S2 cells. The graph to the right depicts the percent undegraded RNA left after 45 min as mean±SE from three repeats.

Notably, immune genes were not significantly up-regulated in the double mutant flies (**Data S1**) and CMTr1 knock-out mice ^34^. Since only a proportion of all mRNAs have cOMe in both *Drosophila* and mice (**Fig. 1e and f**)^9^, the primary role of cOMe is not self/non-self discrimination, at least in *Drosophila*. The relevance of cOMe to prevent detection of non-self RNA by the evolutionary younger vertebrate immune system is linked to the interferon response, which is absent in flies, and they also do not possess unmethylated cap RNA sensors Rig-I and IFITs ^15^.

A potential role of cOMe could be to stabilize mRNA transcripts. However, we find a 3.5 fold increase in up-regulated transcripts compared to down-regulated transcripts in the absence of cOMe, which does not support a general role of cOMe in protecting mRNAs from degradation in *Drosophila*. To further test, whether cOMe protects mRNAs from degradation, we generated fully capped RNA oligonucleotides with or without methylation using the vaccinia capping enzymes and noted that vaccinia CMTr can 2’-*O*-methylate the ribose of the first three nucleotides (fig. S4). When we incubated these RNA oligonucleotides that were uncapped, capped and capped with cOMe in nuclear and cytoplasmic *Drosophila* S2 cell extracts, cOMe did not affect RNA stability, while the lack of a cap resulted in increased degradation, which is consistent with observations in mammalian systems ^35^ (**Fig. 3c**).

### CMTr2 has a dedicated set of target genes

We next investigated how many genes produce mRNAs that contain cOMe. Since the levels of cOMe are low (**Fig. 1e and f**), we reasoned that cOMe is either co-transcriptionally added to mRNAs of only a few specific genes or, of only a fraction of all mRNAs. To distinguish between these two possibilities, we stained polytene chromosomes from larval salivary glands. CMTr1 prominently co-localized with RNA Pol II (**Fig. 4a-e**), suggesting that cOMe is introduced co-transcriptionally and is wide-spread. In contrast, CMTr2 only prominently localized to a subset of transcribed genes suggesting that CMTr2 has a preferred set of target genes (**Fig. 4f-j**).

**Figure 4.**
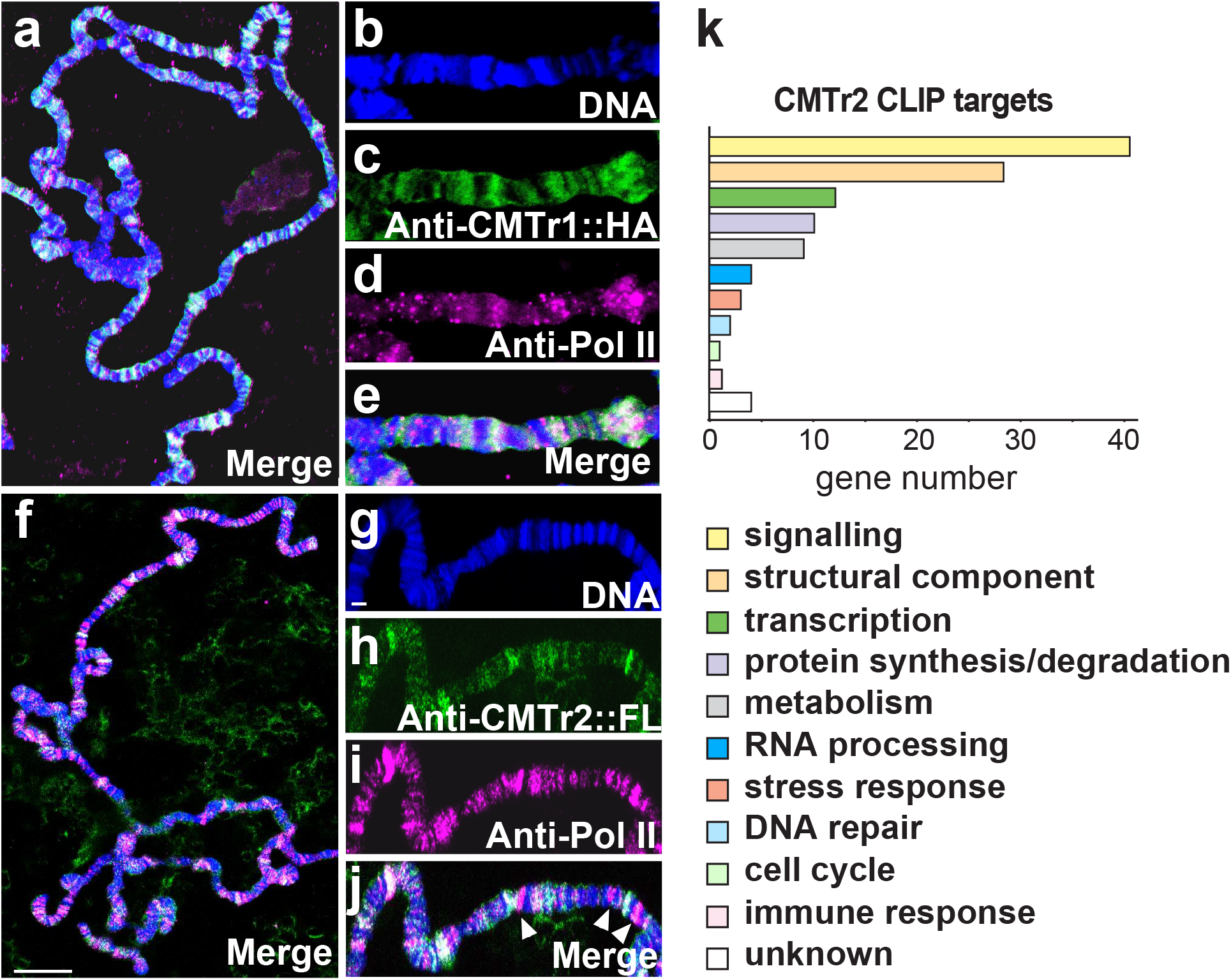
CMTr2 localizes to distinct sites of transcription and has a dedicated set of targets. **(a-l**) Polytene chromosomes from salivary glands expressing CMTr1::HA (a-e) or CMTr2::FLAG (f-j) stained with anti-Pol II (magenta, d), anti-HA (green, c) and DNA (DAPI, blue, b), or merged (white, a and e). Arrow heads indicate absence of CMTr2. Scale bars in f are 10 µm and in g are 2 µm. **(k)** Functional classification of CMTr2 CLIP targets.

We subsequently used CLIP (crosslinking and immunoprecipitation) to identify targets for CMTr1 and CMTr2. For these experiments we used a CMTr double knock-out line which contained genomic rescue constructs for CMTr1 and CMTr2 that are tagged with an HA or FLAG epitope, respectively. From these experiments we obtained 36 and 701 protein coding genes for CMTr1 and CMTr2, respectively, that were twofold or more enriched above input (**Data S3**). Finding so few enriched genes for CMTr1 when CMTr1 co-stained with RNA Pol II on polytene chromosomes indicates that CMTr1 globally associates with most genes. In contrast, the larger number of CMTr2 enriched genes suggests that it introduces cOMe to a more specific set of target transcripts.

To obtain a high confidence catalogue of CMTr2 CLIP targets, we took genes that were at least threefold enriched (117 genes) and analysed them according to gene function. Consistent with previous analysis of differentially expressed genes (**Fig. 3a and b, and Data S1 and S2**), this analysis revealed prominent effects on gene networks involved in cellular signalling including a number of genes encoding ion channels or their regulators and synaptic vesicle release in addition to many cell adhesion molecules (**Fig. 4k, Data S3**). Only few CMTr CLIP targets are differentially expressed in CMTr mutants further supporting that cOMe does not affect mRNA stability (**Data S3)**.

### Cap methylation enhances translation of mRNAs at synapses

Since cOMe can enhance translation in trypanosomes ^36^, we tested whether cOMe is required for local translation at synapses by puromycin incorporation. Indeed, in the absence of CMTrs protein synthesis is significantly reduced at synapses of third instar neuromuscular junctions (**Fig. 5a**).

**Figure 5.**
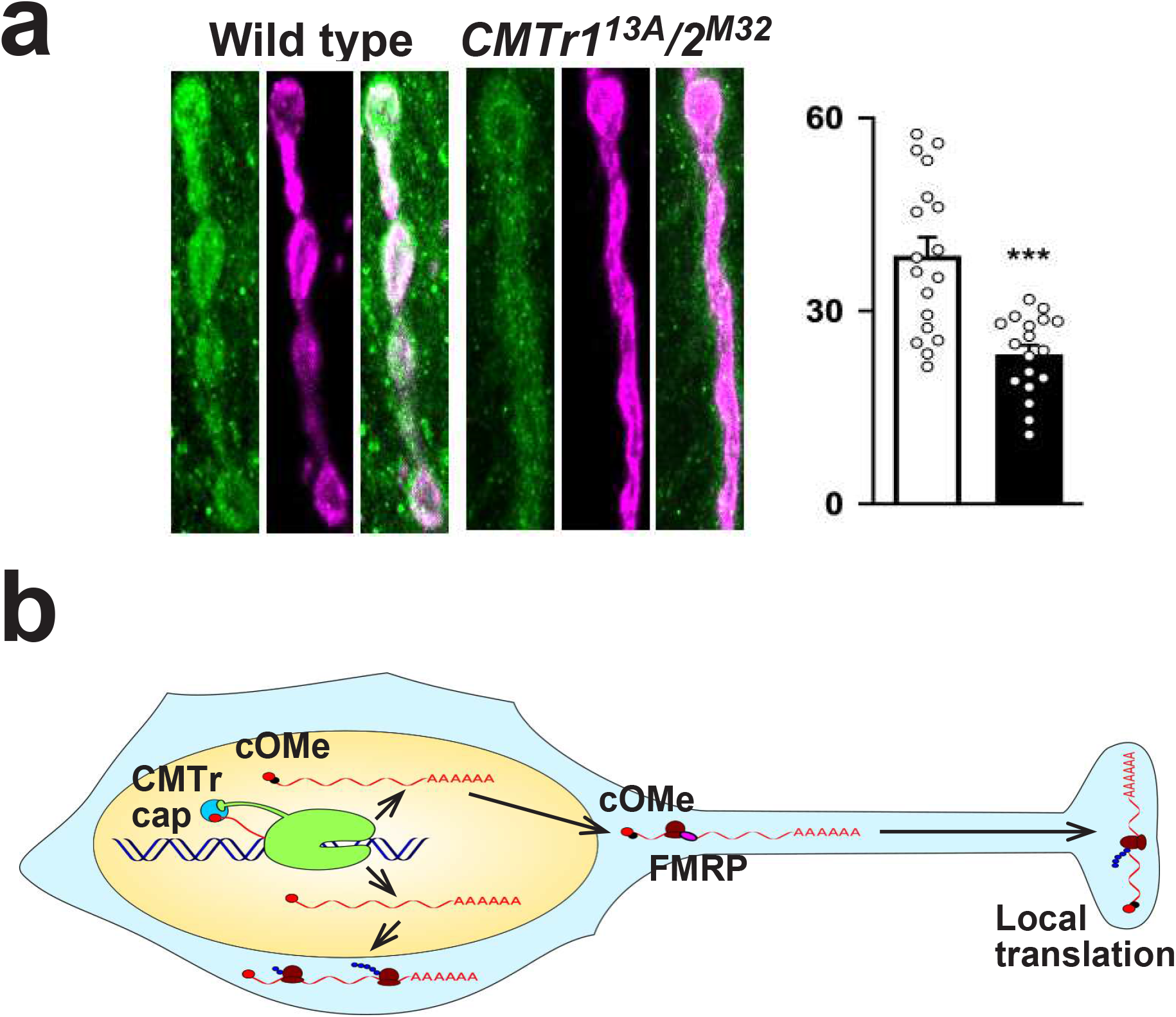
2’-*O*-ribose methylation of mRNA cap-adjacent nucleotides is required for local translation at synapses. **(a)** Staining of synapses at third instar NMJs after puromycin incorporation (green) compared to HRP staining (magenta) in control (left) and *CMTr1*^*13A*^; *CMTr2*^*M32*^ double mutant larvae (right). The mean±SE of the intensity is shown on the right in arbitrary units in white for the control, in black for *CMTr1*^*13A*^; *CMTr2*^*M32*^ double mutant larvae (n=10, ***p<0.0001). **(b)** Model for the role of cap-adjacent 2’-*O*-ribose methylation in gene expression in neurons. FMRP: Fragile X Mental Retardation protein, cOMe: cap 2’-O-ribose methylation at cap-adjacent nucleotides, ribosomes are shown as brown blobs.

## Discussion

Although known for over 40 years, the role of cOMe in animals has been enigmatic due to the lack of knockout models ^15^. Here we show that loss of cOMe has little obvious phenotypic consequences leading to development of healthy and fertile flies. In accordance with prominent expression of mRNA methyltransferases in the brain, however, we find that cOMe is essential for reward learning ^3,4^.

### Tuning protein synthesis in neurons for learning

Short-term reward memory measured immediately after training is considered to be insensitive to blockers of protein synthesis ^29,37^. It therefore seems somewhat enigmatic that cOMe would play an acute role in the reward learning process. Moreover, cOMe occurs in the nucleus before the mRNAs undergo a lengthy journey to the synapse. Our experiments demonstrate a role for cOMe in adult KCs but the two days required to induce CMTr2 expression does not have the required temporal resolution to distinguish between roles before and during learning itself. We therefore currently favor a model for cOMe in establishing/maintaining the appropriate repertoire of locally-translated synaptic factors in adult KC synapses, that are necessary to support reward learning, rather than directly in learning-induced synaptic change. Consistent with prior reports of neuronal localization of mRNAs encoding cytoskeletal proteins, neurotrophins, membrane receptors and regulatory kinases important for synaptic activity and plasticity ^38^, we find that CMTr targets include many cell adhesion and signalling molecules. To mention in particular as CMTr2 target is the *volado*-encoded α-integrin that was shown to be defective in short-term memory performance ^39^. Work in several organisms has also demonstrated roles for neuronal cell adhesion molecules (NCAMs) in acute forms of plasticity and includes *Drosophila* mutants in the N-CAM homolog fasII ^40,41^. Although both of these *Drosophila* studies revealed defects in short-term aversive memory, other locally-translated adhesion molecules could also be specifically required to support short-term and more persistent reward memory.

It is well known that many mRNAs are transported and stored in various cellular locations including dendrites and synapses ^38,42^. In dendrites, translation of mRNAs occurs in polysomes, while in synapses the main form of translation is from monosomes ^43^. Our discovery of a function for mRNA cOMe in learning and local translation of transcripts at synapses (**Fig. 5b**) has important implications in understanding the role of these modifications in affecting gene expression in synaptic plasticity.

## Materials and Methods

### Generation of mutant fly strains

The deletion allele *y w CMTr1*^*13A*^ (excision 13A) and *y w; CMTr2*^*M32*^ (excision M32) were obtained from imprecise excision of transposon *P{EPgy2}CG6379*^*[EY08403]*^ over *Df(X)BSC869* and *P{EP}aft*^*[G6146]*^ over *Df(2R)BSC347* in females and mapped by primers CG6379 F1 (GTCTGGACTTATCGCACCACCTATCG) and R5 Spe (GGTAACTAGTGCTGTGGCCCAACTTGTCCGCAATGAAC), and aft F5 (CCTTCCGAAGTGGAGCAGCTCTTCGAG) and R8 (GGTGGCAGGTAGCATAGTGTCTTGCTTTC). The 192 bp and 287 bp PCR fragments were sequenced for validation. *y w CMTr1*^*13A*^ and *y w; CMTr2*^*M32*^ excision lines were viable when first generated. To normalize genetic backgrounds, excision lines were outcrossed to the Df lines for five generations. A control *y w* line was generated by crossing *Df(X)BSC869* to *P{EP}aft*^*[G6146]*^ and *Df(2R)BSC347* to *P{EPgy2}CG6379*^*[EY08403]*^ for five generations and then combined. To determine survival of mutants, freshly hatched larvae were individually picked and grown in groups of 30 and surviving adults counted.

### Generation of constructs and transgenic fly strains

To clone CMTr1 and CMTr2 cDNAs, total RNA was extracted with Tri-reagent (SIGMA) from larval brains and reverse transcribed with Superscript II as described ^44^. CMTr1 was amplified from this cDNA with primers pUAST CG6379HA F2 (CGAACCTTCGGACGATGAGAACTCGGAGCCCACGCCCAAGAAG) and pUAST CG6379 F3 (GCAGAATTCGAGATCTAAAGAGCCTGCTAAAGCAAAAAAGAAGTCACCATGGACGAACCTTCGGACGATGAGAACTCG) with return primer R5 Spe in a nested PCR with Q5 polymerase (NEB) and cloned with EcoRI and SpeI into a modified pUAST vector containing an attB site for phiC31 mediated integration. The w+-marked *pUAST CMTr1:HA* construct was inserted into attP VK0002 at 76A by phiC31 transgenesis.

CMTr2 was amplified from this cDNA as two fragments with primers aft cDNA F1 (CCTGCTAAAGCAAAAAAGAAGTCACCATGAGCTTTCGTTCGTCTCCGCAGGGAA AGCCAC) and aft cDNA F2 (GGGAATTCGAGATCTAAAGAGCCTGCTAAAGCAAAAAAGAAGTCACCATG) as nested PCR and aft cDNAR2 (CTCATCCTTTTCATATTTGCTATGAAGGTAATGATTCAGAGATGCTATG), and the second fragment with aft cDNA F3 (TACCTTCATAGCAAATATGAAAAGGATGAGATTAAATGGCGCTGGCGCTCAACTACTTTG) and aft cDNA R1 (CTCGGTACCAAATACtGCTGCCGACTCTTGGATGGAACCGACATCTG) with Q5 polymerase (NEB), the two PCR fragments were then fused by PCR and cloned with EcoRI and KpnI into the pUC 3GLA vector ^45^ containing an attB site for phiC31 mediated integration. The GFP+-marked *pUC 3GLA UAS CMTr2:FLAG* construct was inserted into attP40 at 25C by phiC31 transgenesis.

Genomic rescue constructs were made by recombineering from BAC clones. For gCMTr1, the ends were amplified with Q5 polymerase (NEB) using primers dMtr end1F1 (GGCACTAGTgcgcatgaattaagtgctaaaatgtg) and dMtr end1R1 (ATCCCGGCTTATGTGTGTCCAACATG), and dMtr end2F2 (ATCCCAAACCGAACCACATTAAAGG) and dMtr end2R2 (CCGTGGTACCGGTGTTATGCTCGGACAGTGGTAATCGAATG) from BAC DNA prepared as described ^46^ and cloned into pUC 3GLA using SpeI and KpnI. The 10.5 kb genomic fragment was then retrieved using the ends vector linearized with EcoRV from BacR21I10 as described ^45^. The C-terminal HA TEV myc tag was then incorporated by PCR into a 495 bp AvrII and SbfI fragment and cloned with these sites. The GFP+-marked *pUC 3GLA gCMTr1:HATEVmyc* construct was inserted into attP VK0022 at 57F by phiC31 transgenesis. For CMTr2, the ends were amplified with Q5 polymerase (NEB) using primers aft end1 F1 Bam (CCAGGATCCGCGGCCGCATGGGAGGTATGCGATTAATGGC) and aft end1 R1 Xba (CCTCTAGAGGCCTAAATTTGAAATAGTTATCTCCATATAATATTTATGAG), and aft end2 F2 Xba (GCCTCTAGAGGCCTGTTTCTCACCCATTACGC) and aft end2 R2 PvuII (CTGATCCCTGGAAGTAAAGATTCTCGGTACCAAATACTGCTGCCGACTCTTGGATGGAAC) from BAC DNA and cloned together with a linker BirA FLAG linkA (CTGGAGGATTAAATGACATCTTTGAAGCACAGAAGATCGAATGGCATGAGGATTACAAGGACGACGATGACAAGGCTTGA) and BirA FLAG linkB (CTAGTCAAGCCTTGTCATCGTCGTCCTTGTAATCCTCATGCCATTCGATCTTCTGTGCTTCAAAGATGTCATTTAATCCTCCAG) into a modified pUAST using BamHI and SpeI in a four-way ligation. The 6.7 kb genomic fragment was then retrieved using the ends vector linearized with StuI from BacR20E20 as described ^45^. The w+-marked *CASPR gCMTr2: TEVFLAG* construct was inserted into attP40 at 25C by phiC31 transgenesis.

Essential parts of all DNA constructs were sequence verified.

### Behavioral assays

For negative geotaxis experiments, groups of 20 flies kept in two inverted fly vials (19 cm) were tapped to the bottom. A movie was then made to record the moving flies and a frame about 5 sec after the flies started running upwards and before the first fly reached the top was taken to measure the distance the flies have run upwards.

For learning and memory experiments, two to five day old flies of both sexes were used for behavioral experiments in a T-maze. Odors used were 4-methylcyclohexanol (MCH) and 3-octanol (OCT).

For appetitive learning and 24 hour memory testing, flies were starved for 21–23 h prior to training and training was done as described ^29^. Briefly, a group of about 120 flies were exposed first to the unconditioned odor (CS−) for two minutes followed by 30 seconds of air, and then to the conditioned odor (CS+) in the presence of dry sucrose for two minutes. For appetitive learning or immediate memory, flies were tested immediately after training for their choice between the two odors. For 24 hour memory, flies were transferred into a standard cornmeal food vial after training and after one hour, they were transferred into food-deprivation vials until testing on the next day.

Odor and sugar acuity tests were performed as described in ^47^ with some modifications. For odor acuity tests, starved flies were directly placed into the T-maze to test for odor avoidance (OCT or MCH) against the smell of plain mineral oil. For sugar acuity test, a filter paper with size 18×8cm was placed into a glass milk bottle (250ml). Half of the filter paper (∼9×8cm) was soaked with saturated sucrose and dried before use. For the test, starved flies were placed into the bottle and the number of flies on both parts of the filter paper were counted separately two minutes later. The performance index was calculated as [N_sugar_/N_total_] × 100, where N_total_ = N_sugar_ + N_plain_.

For conditional expression GSG GAL4 was used ^48^, that is activated by feeding flies with the progestin, mifepristone (RU486). Accordingly, flies were kept on RU486 (200 µM (SIGMA), 5% ethanol) or control (5% ethanol) standard fly food for two days at 18°C before starvation and training.

### Statistical analysis of behavioral data

Behavioral data was analyzed using GraphPad Prism 6. Two-tailed t tests were used for comparing two groups, and one-way ANOVA followed by a Tukey’s post-hoc test was used for comparing multiple groups.

### Analysis of cap-adjacent 2’-*O*-ribose methylation

Total RNA was extracted with Trizol (Invitrogen) and PolyA mRNA from two rounds of oligo dT selection was prepared according to the manufacturer (Promega). Alternatively, polyA mRNA from one round of oligo dT selection was followed by ribosomal RNA depletion using biotinylated oligos as described ^49^. For each sample, 50 ng of mRNA was decapped using either tobacco acid pyrophosphate (250 U; Epicenter) or RppH (NEB) in buffer provided by the supplier and then dephosphorylated by Antarctic phosphatase (NEB). The 5’-end of dephosphorylated mRNAs were then labeled using 10 units of T4 PNK (NEB) and 0.5 µl [γ-^32^P] ATP (6000 Ci/mmol, 25 µM; Perkin-Elmer). The labeled RNA was precipitated, and resuspended in 10 µl of 50 mM sodium acetate buffer (pH 5.5) and digested with P1 nuclease (SIGMA) for 1 h at 37° C. Two microliters of each sample was loaded on cellulose F TLC plates (20×20 cm; Merck) and run in a solvent system of isobutyric acid:0.5 M NH_4_OH (5:3, v/v), as first dimension, and isopropanol:HCl:water (70:15:15, v/v/v), as the second dimension. TLCs were repeated from biological replicates. The identity of the nucleotide spots was determined as described ^9,50^. For the quantification of spot intensities on TLCs, a storage phosphor screen (K-Screen; Kodak) and Molecular Imager FX in combination with QuantityOne software (BioRad) were used.

For the analysis of CAGEseq data, nucleotides in the N1 position of mRNA following the m^7^G artifact were counted in a loop using grep in bash on all fastq files available from SRP131270 (data GSE109588) ^24^. Counting lines with the pattern ‘…CAGCAGGN……….’ where N was replaced with the nucleotide being counted. Similarly, nucleotides were counted in the m^7^G artifact position using grep with the pattern ‘…CAGCAGN ’ and in the second position following the m^7^G artifact using ‘…CAGCAGG.N ’.

### Generation of S2 cell extracts

S2 cells (ATCC) were cultured in Insect Express medium (Lonza) with 10% heat-inactivated FBS and 1% penicillin/streptomycin. Extracts were made after the Dignam protocol with modifications ^51,52^. Cells were washed in PBS and resuspended in five times the packed cell volume in buffer A (15 mM HEPES, pH 7.6, 10 mM KCl, 5 mM MgCl_2_, 350 mM sucrose, 0.1 mM EDTA, 0.5 mM EGTA, 1 mM DTT, 1 mM PMSF (stock 0.2 M in isopropanol), 1 µg/ml leupeptin), spun down with 3000 g for 5 min and resuspended in buffer A and allowed to swell for 10 min on ice. Cells were then homogenized with a Dounce homogenizer with the loose pestle (B) with approximatively 15 up and down strokes until cells were 80-90% lysed. The extract was then spun at 4000 g for 15 min, the supernatant taken off and 0.11 volume buffer B (10x: 0.3 M HEPES, pH 7.6, 1.4 M KCl, 30 mM MgCl_2_,) added. The supernatant was then spun at 34 000 g for 1 hour. The resulting supernatant is the S-100 cytoplasmic extract. The nuclei were the resuspended in 50% of the volume in buffer C (20 mM HEPES, pH 7.6, 420 mM NaCl, 1.5 mM MgCl_2_, 0.2 mM EDTA, 0.5 mM DTT, 1 mM PMSF (stock: 0.2M in isopropanol), 1 µg/ml leupeptin, 25 % v/v glycerol (Ultrapure, Gibco) using a pipette, a stirrer added, the volume slowly increased by another 50% of the nuclei volume with buffer C and then the nuclei were extracted for 30 min. The extract was then spun 30 min at 10 000 g at 4° C and the supernatant taken off without the the white slur on top. This extract was then dialyzed in buffer E (20 mM HEPES, pH 7.6, 100 mM KCl, 0.2 mM EDTA, 0.5 mM DTT, 1 mM PMSF (stock: 0.2M in isopropanol), 1 µg/ml leupeptin, 20 % v/v glycerol (Ultrapure, Gibco) for 2 hours. After dialysis, the supernatant was spun at 10 000 g for 10 min and aliquots frozen in liquid nitrogen and extracts stored at -80° C.

### Generation of a cap labelled probe, UV-crosslinking, RNA stability assay, immunoprecipitation and denaturing gel electrophoresis

As probe for UV-crosslinking, RNA stability and binding experiments the trypanosome splice leader oligo (trypSL, AACUAACGCUAUUAUUAGAAC)^53^ was used. 6 pmole trypSL (1.25 µl from a 50µM stock was kinased with 2 µl ^32^PgammaATP (25 µM, 6000Ci/mmol, 150 mCi/ml, Perkin Elmer) with 10 U PNK in 10 µl with 20 U RNasin (Roche). After 1 h, the probe was extracted by phenol/CHCl_3_ and precipitated. The second phosphate was then added with Myokinase (Sigma M3003, Myokinase was dialyzed into 100 mM NaCl, 50 mM TrisHCl pH 7.5, 1 mM MgCl_2_, 1 mM DTT), 100 U in 20 µl, in a total volume of 40 µl to 2.4 pmole trypSL in the presence of 1 mM ATP and 20 U RNasin (Roche) in vaccinia capping buffer. After 2 h, the RNA was extracted by phenol/CHCl_3_ and precipitated. Capping was then done in 20 µl with vaccinia capping enzymes (NEB) according to the manufactures instructions and after 90 min 2 µl terminator nuclease buffer A and 0.7 U Terminator nuclease (Epicenter) were added. After 30 min, the RNA was extracted by phenol/CHCl_3_ and precipitated. The RNA was then analysed on 20% polyacrylamide gels, dried and exposed to a phosphoimager screen. RNAse I digestion to analyse 2’-O-ribose methylation was done in the presence of 10 U T4 PNK (NEB) in 50 mM Tris-acetate (pH 6.5), 50 mM NaCl, 10 mM MgCl_2_ and 2 mM DTT to remove 2’,3’-cyclic phosphate intermediates ^54^.

UV-crosslinking was done as described ^52^. Briefly, ^32^P labeled capped trypanosome splice leader oligo with or without cOMe was incubated in a total volume of 10 µL, in 40% (v/v) nuclear or cytoplasmic extract, 1 mM ATP, 5 mM creatine phosphate, 2 mM MgAcetate, 20 mM KGlutamate, 1 mM, DTT, 20 U RNasin (Roche), and 5 µg/mL tRNA at room temperature for 25 min and UV cross-linked on ice at 254nm for 20 min in a Stratalinker (Stratagene), followed by digestion with RNase A/T1 mix (Ambion) at room temperature for 15 min. Samples were then taken up in SDS-protein gel buffer and run on 8% gels, the gels dried and exposed to a phosphoimager screen.

For RNA stability experiments, ^32^P labeled uncapped and capped trypanosome splice leader oligo with or without cOMe was incubated in a total volume of 10 µl, in 40% (v/v) nuclear or cytoplasmic extract, 1 mM ATP, 5 mM creatine phosphate, 2 mM MgAcetate, 20 mM KGlutamate, 1 mM, DTT, 20 U RNasin (Roche), and 5 µg/mL tRNA on ice for 45 min. Input was take before the addition of nuclear extract. The RNA was extracted by phenol/CHCl_3_ and precipitated. Samples were then separated on 8% polyacrylamide gels, dried and exposed to a phosphoimager screen.

For immunoprecipitations of CBP80, ^32^P labeled capped Trypanosome splice leader oligo with or without cOMe was incubated in a final volume of 120 µl in IP-Buffer (150 mM NaCl, 50 mM Tris HCL, pH 7.5, 1% NP-40, 5% glycerol) together with nuclear extract (40%, v/v), rabbit anti-CBP80 (4 µl, gift from D. Kopytova), 20 µl protein A/G beads (SantaCruz) in the presence of Complete Protein Inhibitor (Roche) and 40 U RNase inhibitors (Roche) for 2h at 4° C. After washing the beads, RNA was extracted by phenol/CHCl_3_ and precipitated. Samples were then separated on 8% polyacrylamide gels, dried and exposed to a phosphoimager screen.

### Immunostaining of tissues

In situ antibody stainings were done as described previously ^31^ using rat anti-HA (MAb 3F10, 1:20; Roche), rabbit anti-FLAG (M2, 1:250, SIGMA), mouse anti-ELAV (MAb 7D, 1:20, which recognizes 7 amino acids unique to ELAV) and anti-GFP (1:250; Invitrogen A11122) and visualized with Alexa Fluor 488- and/or Alexa Fluor 647-coupled secondary antibodies (1:250; Molecular Probes or Invitrogen, A11034). DAPI (4’,6-diamidino-2-phenylindole) was used at 1 µg/ml. For imaging, tissues were mounted in Vectashield (Vector Labs) for confocal microscopy using a Leica TCS SP5/SP2. Images were processed using Fiji.

To analyse synapses at neuromuscular junctions (NMJ) third instar wandering larvae were dissected in PBS and fixed with Bouin’s solution (Sigma-Aldrich, HT10132) for 5 minutes using. The samples were washed three times in PBT (PBS with 0.1% Triton™ X-100 (Sigma, T8787) and 0.2% BSA) for 15 minutes. Primary antibody were rat anti-HA (MAb 3F10 1:20, Roche) or rabbit anti-FLAG (M2, 1:250, SIGMA), Mouse anti-NC82 (1:50, DSHB), rabbit anti-CBP80 (1:100, Gift from D. Kopytova)^55^ and DAPI (4=,6=-diamidino-2-phenylindole,1 µg/ml) was carried out overnight at 4° C followed by secondary antibodies (conjugated with Alexa Fluor 488 or Alexa Fluor 647 (1:250; Molecular Probes, Invitrogen) at RT for 4-5 hours. NMJs were mounted in Vectashield (Vector Labs), scanned with Lecia TCS SP8 and processed using FIJI. For quantification of synapse stainings the mean intensity of the boutons was calculated using the Nikon NIS-Elements Basic Research (BR) imagining software, and the data was analysed using GraphPad Prism.

### Polytene chromosome preparations and stainings

CMTr1 and CMTr2 were expressed in salivary glands with *elav*^*C155*^*-GAL4* from a *UAS* transgenes tagged with HA or FLAG, respectively, as described ^33^. Briefly, larvae were grown at 18° C under non-crowded conditions. Salivary glands were dissected in PBS containing 4% formaldehyde and 1% TritonX100, and fixed for 5 min, and then for another 2 min in 50% acetic acid containing 4% formaldehyde, before placing them in lactoacetic acid (lactic acid:water:acetic acid, 1:2:3). Chromosomes were then spread under a siliconized cover slip and the cover slip removed after freezing. Chromosome were blocked in PBT containing 0.2% BSA and 5% goat serum and sequentially incubated with primary antibodies (mouse anti-PolII H5 IgM, 1:1000, Abcam, and rat anti-HA MAb 3F10, 1:50, Roche, or rabbit anti-FLAG, 1:1000, SIGMA) followed by incubation with Alexa488- and/or Alexa647-coupled secondary antibodies (Molecular Probes) including DAPI (1 µg/ml, Sigma).

### Illumina sequencing and analysis of differential gene expression

For sequencing, QuantSeq 3’ FWD libraries were generated from *y w* control and *y w CMTr*^*13A*^; *CMTr*^*M32*^ flies. The QuantSeq 3’ FWD kit was used according to the manufacturer’s instructions with the following modifications: RNA was not denatured, and 6 U of Heparinase I (NEB) was added to the first strand cDNA synthesis mix. Pooled indexed libraries were sequenced on an Illumina NextSeq 500 to yield between 10 and 30 million single-end 50bp reads per sample.

After demultiplexing with Illumina bcl2fastq v1.8.4, sequence reads were aligned to the Drosophila genome (dmel r6.02) using STAR 2.6. Reads for each gene were counted using HTSeq-count and differential gene expression determined with DESeq2 and the Benjamini-Hochberg for multiple testing to raw P-values with p<0.05 considered significant.

### CLIP of CMTr targets

For CLIP, RNA was prepared essentially as described from 14-18h old embryos of *y w CMTr*^*13A*^; *gCMTr1:TEVHA CMTr*^*M32*^ *gCMTr2:TEVFLAG* ^56^. Embryos were first dechorionated in 50% bleach, washed and then fixed in heptane containing 5% formaldehyde (10 ml heptane, 1.75 ml 37% formaldehyde, and 1.3 ml PBS equilibrated for 30 min) for 10 min with vigorous shaking. Embryo extracts were then prepared in RIPA buffer (150 mM NaCl, 50 mM Tris–HCL, pH 7.5, 1% NP-40, 0.5% Na-deoxycholate, 0.05% SDS) in a 1-ml Dounce homogenizer. After 20–40 strokes with the tight pestil, 1 vol of immuno-precipitation (IP) buffer was added (150 mM NaCl, 50 mM Tris–HCL, pH 7.5, 0.05% NP-40). The extract was then cleared by centrifugation for 15 sec. IPs were done with monoclonal anti-HA antibodies coupled beads (Sigma) or anti-FLAG antibodies and protein A/G beads (SantaCruz) in IP buffer containing 7 mM CaCl_2_, 40 U of RNase inhibitor (Roche), 2 U of TurboDNase (Ambion), and 15% of extract for 2 hr at room temperature. After washing and TEV Proteinase (Promega) digestion for 1 h on ice, the supernatant was taken off, Proteinase K digested (0.5 mg/ml in 150 mM NaCl, 100 mM Tris–HCl, pH 7.5, 10 mM EDTA, 0.25% SDS) for 30 min at 37° C, and RNA was isolated by phenol/chloroform extraction and ethanol precipitation in the presence of glycogen.

The RNA was then reverse-transcribed with Superscript II (Invitrogen) according to the manufacturer’s instructions using a random nonamer tagged partial p7 sequence (CACGACGCTCTTCCGATCTNNNNNNNNN) and the first strand synthesis product was purified using AMPure XP beads (Beckman) following the manufacturer’s instructions with 1.8 volumes. To generate double stranded cDNA and sequencing-ready libraries, Lexogen’s quant-seq 3’FWD kit was used, proceeding from the RNA removal and second strand synthesis steps. The input library was generated the same way from RNA before IP. Size selection of libraries were carried out with PAGE prior to sequencing with the NextSeq 500. Differential gene expression analysis performed as above then provided a simple route to detecting enriched transcripts following immuno-precipitation.

## Data analysis

GO enrichment analysis was performed with Pantherdb. Gene expression data were obtained from flybase. Visualization of RNA-seq data were carried out with the R packages EnhancedVolcano Version 1.4.0 and ggplot2 in the R studio environment ^57,58^.

Hypergeometric p-values for the significance of overlapping genes between CMTr2 and FMRF CLIP targets were calculated using the 197 successes in a sample size of 701 CMTR2 clip targets, compared to the 2432 successes of FMRP targets in cholinergic and GABAergic neurons ^59^ in the whole population of 17421 coding genes that can be returned following alignment (p=1.34^-23^).

## Data availability

All data are available in the main text or the supplementary material and gene expression data have been deposited at GEO under the accession numbers GSE116212 and GSE138868.

## Acknowledgments

We thank S. Brogna, C. Samakovlis, B. Suter, J-Y. Roignant, M. Ramaswami, FlyORF and the Bloomington and Kyoto stock centers for fly lines, S. Brogna, D. Kopytova, P. Lasko and the Developmental Studies Hybridoma Bank for antibodies, the University of Cambridge Department of Genetics Fly Facility for injections, BacPAc for DNA clones, E. Zaharieva for help with stainings, D. Balacco and F. Stappers for help with art work, and B. Muller and R. Michell for comments on the manuscript. MS is funded by the BBSRC (BB/R002932/1) and the Leverhulme Trust, RGF from BBSRC (BB/R001715/1), DH from WISB, a BBSRC/EPSRC Synthetic Biology Research Centre (BB/M017982/1) and BBSRC (BB/L006340/1), and S.W. by a Wellcome Principal Research Fellowship (200846/Z/16/Z) and an ERC Advanced Grant (789274). NA is a Nottingham Research Fellow funded by the University of Nottingham.

## Author contributions

IUH and MS performed biochemistry, molecular biology and genetic experiments, YW and SW performed learning experiments and anatomical analysis of adult brains, MPN performed antibody stainings, NA, ZB and RF performed sequencing and biochemistry experiments. NA and DH analyzed sequencing data. IUH and MS conceived the project and wrote the original draft of the manuscript. S.W, R.F., Z.B., N.A., D.H. and all other authors reviewed and edited. M.S., S.W. R.F. and D.H. supervised and acquired funding.

## Competing interests

The authors declare no competing interests.

## Notes

### Competing Interest Statement

The authors have declared no competing interest.

